# *Streptococcus pneumoniae* serotypes carried by young children and their association with Acute Otitis Media during the period 2016 – 2019

**DOI:** 10.1101/2020.11.16.386052

**Authors:** Esra Ekinci, Stefanie Desmet, Liesbet Van Heirstraeten, Colette Mertens, Ine Wouters, Philippe Beutels, Jan Verhaegen, Surbhi Malhotra-Kumar, Heidi Theeten, NPcarriage group

## Abstract

**Background:** *Streptococcus pneumoniae* (Sp) is a major cause of acute otitis media (AOM). Pneumococcal conjugate vaccine (PCV) programs have altered pneumococcal serotype epidemiology in disease and carriage. To establish the clinical picture of AOM in young children exposed to the PCV program in Belgium and the Sp strains they carry, a cross-sectional study started in 2016.

**Material/methods:** In three collection periods from February 2016 to May 2018, nasopharyngeal swabs and background characteristics were collected from children aged 6-30 months either presenting at their physician with AOM (AOM-group) or healthy and attending day care (DCC-group). Sp was detected, quantified, and characterized using both conventional culture and qPCR. Clinical signs of AOM episodes and treatment were registered by the physicians.

**Results:** Among 3264 collected samples, overall pneumococcal carriage and density were similar in AOM (79.2% and 0.50 ×10^6^ copies/μl) and DCC (77.5% and 0.42 ×10^6^ copies/μl). Non-vaccine serotypes were most frequent: 23B (AOM: 12.3%; DCC: 17.4%), 11A (AOM: 7.5%; DCC: 7.4%) and 15B (AOM: 7.5%; DCC: 7.1%). Serotypes 3, 6C, 7B, 9N, 12F, 17F and 29 were more frequent in AOM than in DCC, whereas 23A and 23B frequencies were lower. Antibiotic susceptibility of Sp strains was similar in both groups . No predictors of AOM severity were identified, and 77.3% received an antibiotic prescription.

**Conclusion:** Young children with AOM did not carry Sp more frequently or at higher load than healthy children in day care, but some ST were more frequent in AOM and are not included in the currently used vaccines.

## Introduction

Acute otitis media (AOM) is one of the most common paediatric infections and has been described as one of the leading causes of antibiotic prescription in children in industrialized countries [1, 2]. It is estimated that 80-90% of children have an episode of acute otitis media (AOM) before the age of three, with a peak incidence between six and fifteen months [3, 4].

Streptococcus pneumoniae (Sp) is the most common OM pathogen and has been associated with first or early otitis media episodes, more severe AOM signs and symptoms, and potential middle-ear damage [5]. It colonizes the nasopharynx within the first few weeks or months of life, although carriage is a highly dynamic process [6, 7]. Sp is however commonly a quiescent colonizer of the upper respiratory tract (URT) that only becomes pathogenic when changes in the upper respiratory tract allow the organism to achieve a pathogenic inoculum that overcomes innate and adaptive host defence. Capacity to colonize and invade, differs and it is related to the capsular serotype as well as to the history of colonization with this serotype. So far, at least 100 distinct capsular serotypes have been described [8, 9]. Inactivated pneumococcal conjugate vaccines (PCV) were developed to prevent invasive disease caused by the most pathogenic serotypes. In addition, administration of PCV is associated with a modest reduction of AOM incidence [3]. Before PCV introduction, the most common serotypes causing AOM worldwide were 3, 6A, 6B, 9V, 14, 19A, 19F and 23F [10–13].

Since PCV implementation in children, a tremendous decrease in vaccine-type related carriage and invasive pneumococcal diseases cases has been observed in countries with a high vaccine uptake [14]. However PCVs only target a limited number of (frequently invasive) serotypes and the impact on AOM is less than for invasive diseases. Furthermore, the protection against the vaccine serotypes opens new nasopharyngeal niches for colonization, which favours conditions for serotype replacement. Hence, PCV introduction induced worldwide significant changes in the epidemiology of Sp in disease, carriage and antimicrobial (non-)susceptibility. A French study, evaluating the changes in serotype distribution in children with AOM between 2001 and 2011, showed a significant decrease in PCV7 serotypes. However, there was an increase in the six additional serotypes included in PCV13 and the non-vaccine serotypes 15A and 23A [10]. In another study investigating the pneumococcal serotype distribution in children with AOM in Spain, serotypes 19A and 3 were found as the most common serotypes in the late PCV7 era. The most common serotypes in the early PCV13 era were serotypes 19A and 15C [15]. In USA, an immediate decrease in carriage of serotypes 6A and 19A was observed in healthy children after licensure of PCV13, however non-vaccine serotypes 16, 21 and 35B increased [16]. Belgium has a unique pneumococcal vaccine program history: serotype coverage moved from PCV7 to PCV13 (in 2011) to PCV10 (in 2016), and only very recently (in 2019) the switch back to PCV13 was made. After the PCV13-to-PCV10 switch, the carriage proportion of PCV13-non-PCV10 serotype 19A showed an increasing trend in healthy children in day care centres (DCC)[2]. No recent data were available on the serotype distribution and antibiotic susceptibility of pneumococcal isolates in Belgian children with AOM, and their relationship with clinical symptoms. The current cross-sectional study investigates nasopharyngeal carriage of Sp serotypes in infants with AOM and compares them with healthy infants attending DCC in Belgium.

## Material and methods

The data were collected in an ongoing study to monitor nasopharyngeal carriage in infants in Belgium since 2016. In the first 3 years, this study investigated nasopharyngeal carriage in two infant populations with high reported carriage of Sp, namely healthy infants attending DCC and infants suffering from AOM. The study protocol was described in detail by Wouters, I et al [17, 18] and is summarized here. The study was in line with the Declaration of Helsinki, as revised in 2013. Approval to conduct the current study with ID 15/45/471 was obtained from the University of Antwerp and University Hospital of Antwerp ethics committee (Commissie voor Medische Ethiek van UZA/UA) on 30 November 2015.

The recruitment was restricted to non-summer seasons (November to end May) from 2016 to 2018. In period 1, sampling was performed between January and June 2016, i.e. during and shortly after the switch from PCV13 to PCV10 in the Belgian vaccination program, while in period 2 (2016-2017) and period 3 (2017-2018) samples were collected between November and May. Randomly selected DCCs and hospitals/ paediatric outpatient services were located over the three Belgian regions (the Flemish Region, the Walloon Region and the Brussels-Capital Region). Age limits for inclusion in both populations were 6 months and 30 months. In the first recruitment season, region-specific age criteria were added to include only infants who were offered PCV13 for both primary vaccine doses in the first year of life. AOM was defined by the acute onset of symptoms i.e. within the preceding seven days and specified clinical inclusion (otoscopic) criteria, such as otalgia, middle ear effusion, fever, redness and bulging of the tympanic membrane or perforation. Exclusion criteria for AOM infants as well as healthy infants in DCC were 1) use of antibiotics in the past seven days, 2) presence of a chronic and severe associated pathology and 3) previous inclusion in the study within the same collection period. AOM infants who were recruited in hospitals (outpatient services) had extra exclusion criteria: 1) referral by a GP or 2) having received three or more antibiotic treatments in the past three months.

### Nasopharyngeal samples (NP) and questionnaire

Parents consenting to participate received a questionnaire regarding the infants demographics and clinical characteristics, and pneumococcal vaccination status. A trained nurse or a general practitioner/paediatrician collected a single nasopharyngeal (NP) sample with a flocked nylon swab in 1mL STGG (skim milk, tryptone, glucose and glycerol). Signs of common cold in DCC infants were defined as coughing and/or running nose at sampling.

The samples were stored frozen and assessed by culture at the Reference Centre for Pneumococci (University Hospitals in Leuven) and PCR analysis at the Laboratory of Medical Microbiology (University of Antwerp). NP samples were cultured, Sp strains were serotyped by performing Quellung-reaction and antimicrobial susceptibility to penicillin, tetracycline, erythromycin, levofloxacin and cotrimoxazole (the most frequently used antibiotics in respiratory disease over all ages) was determined by disc diffusion following CLSI 2016 guidelines. If non-susceptibility for levofloxacin or penicillin was identified, the minimal inhibitory concentration (MIC) was determined. A MIC of >2 mg/L and >0.06 mg/L was interpreted as levofloxacin and penicillin non-susceptible, respectively. Quantitative Taqman real-time PCR (qRT-PCR) targeting *lytA*, using StepOnePlus™ Real-Time PCR System (Applied Biosystems ™) was performed to screen the samples for presence of Sp. Bacterial densities were determined based on a standard curve that was set up using 10-fold serially diluted *lytA* PCR product of Sp strain ATCC 49619. Samples and standard curves were run in triplicate. Samples were considered positive for Sp if either culture or PCR (cut off cycle threshold value ≤35.0) was Sp-positive [17]. *Lyt*A positive samples were pooled and screened for presence of PCV13-non-PCV10 serotypes (3, 6A, 19A) [2].

### Clinical score for AOM severity

A clinical severity score was developed based on a clinical evaluation by the treating physician. Physicians were classified as prescriber (prescribed antibiotics in >50% of AOM episodes included in the study) and non-prescriber (prescribed antibiotics in ≤50% of AOM episodes). Parameters included in the score were temperature (<38°C=0; 38-38.5°C=1; 38.6-39°C=2 ; >39°C=3), otorrhea (none=0; unilateral=1; bilateral=2), middle ear effusion (none=0; unilateral=1; bilateral=2), bulging of the tympanic membrane (none=0; unilateral=1; bilateral=2) and prescription of antibiotic medication (no=0; by a prescribing physician=1; by a non-prescribing physician=2). The sum of these scores comprised the total clinical score (0-11). If one parameter was missing, the sum of the remaining scores was converted to a denominator of eleven. In 6 children in whom two or more parameters were missing, the clinical score was not determined.

### Statistical analyses

All descriptives and statistical analyses were calculated in JMP Pro 14.3. To test for significant differences in proportions the Chi-Square Test (Chi2), or Fisher’s Exact Test (FET) where appropriate, were used. To compare overall association between vaccination status and health status, bivariate logistic regression was performed using period and vaccination status as factors and AOM/DCC as outcome. To compare parameters that were not normally distributed the Mann-Whitney U Test (MWU) or Kruskal-Wallis Test (KW) were used as appropriate.

The sample size of the main study was calculated according with its primary aim to compare carriage prevalence of three Sp vaccine serotypes over the first three years (2016-2018). For the current study comparing nasopharyngeal carriage of Sp in infants with AOM and in infants attending DCC, a post hoc power analysis was performed using R package power.

## Results

### Recruitment and infant characteristics

Over the three-year study period, a total of 3264 nasopharyngeal samples were collected of which 3175 were included in the analysis (table 1). 74 nasopharyngeal samples from DCC children and fifteen nasopharyngeal samples from AOM children were excluded due to use of oral antibiotics in the seven days prior to sampling, inappropriate age, presence of a severe associated pathology or because of a missing informed consent. Moreover, four children with AOM were excluded from the clinical analysis regarding AOM symptoms, due to incomplete questionnaires. A random selection of 194 DCC-samples collected in 2016-2017 was analysed only by culture and not by PCR due to restriction at laboratory level.

**Table 1:** Number of included samples, DCCs and clinics per sampling period.

|  | **Period 1**<br>**2016** | **Period 2**<br>**2016 – 2017** | **Period 3**<br>**2017 – 2018** |
| --- | --- | --- | --- |
| **DCC setting** | 85 | 112 | 102 |
| **DCC samples** | 769 | 1096 | 953 |
| **AOM setting** | 12 | 21 | 19 |
| **AOM samples** | 39 | 122 | 205 |

A post hoc power calculation was performed taking the DCC (N=2809) and AOM (N=366) sample size into account. The sample size allows for the detection of a 15% difference in carriage prevalence of Sp-serotypes between healthy DCC infants and AOM-infants with 77% power at a significance level of 5% (two-sided).

AOM children differed in several demographic and clinical characteristics from DCC children (Table 2). Overall, children with AOM tended to be younger than children recruited in DCCs. The convenience recruitment of AOM children resulted in a disproportionate regional distribution, which was not the case for DCC children. Overall, breastfeeding for more than six months was more frequent in DCC children and parental smoking was more frequent in AOM children. Children with AOM more often had a history of previous hospitalisation, in comparison to DCC children, and more frequently had siblings (DCC: 62.5%; AOM: 67.4%). A significantly higher proportion of AOM children had a history of one or more previous AOM episodes over the three-year study period (AOM-history in DCC: 26.9%; in AOM: 64.3%) compared to DCC children. Lastly, antibiotic use in the past three months was more frequent in DCC children than AOM children; but it has to be noted that only in the latter the number of antibiotic treatments was an exclusion criterium (see methods section).

**Table 2:** Overall demographic and clinical characteristics of children with AOM and children attending DCC.

|  |  | **Children with AOM**<br>**N=366** |  | **Children attending DCC**<br>**N=2809** |  | **Statistical significance**<br>**(p-value for**<br>**AOM vs DCC)** |
| --- | --- | --- | --- | --- | --- | --- |
|  |  | **n** | **%** | **n** | **%** |  |
| **Region** | Flanders | 197 | 53.8 | 1489 | 53.0 | **0.007** |
|  | Wallonia | 141 | 38.5 | 950 | 33.8 |  |
|  | Brussels | 28 | 7.7 | 370 | 13.2 |  |
| **Age (months)** | 6-12 | 163 | 45.0 | 578 | 20.6 | **<0.001** |
|  | 13-24 | 154 | 42.5 | 1497 | 53.3 |  |
|  | 25-30 | 45 | 12.4 | 734 | 26.1 |  |
| **Gender** | male | 186 | 51.4 | 1421 | 50.6 | 0.786 |
| **Preterm delivery** | yes | 25 | 6.9 | 223 | 8.0 | 0.480 |
| **Breastfeeding >6 months** | yes | 81 | 22.5 | 927 | 33.1 | **<0.001** |
| **At least one parent smokes** | yes | 103 | 28.6 | 586 | 21.0 | **<0.001** |
| **Previous hospitalisation** | yes | 108 | 30.0 | 514 | 18.5 | **<0.001** |
| **Number of siblings** | 0 | 117 | 32.6 | 1038 | 37.5 | **<0.001** |
|  | 1 | 121 | 33.7 | 1242 | 44.9 |  |
|  | >1 | 121 | 33.7 | 488 | 17.6 |  |
| **AOM-history** | 0 | 101 | 35.7 | 1994 | 73.1 | **<0.001** |
|  | 1 | 86 | 30.4 | 279 | 10.2 |  |
|  | >1 | 96 | 33.9 | 454 | 16.7 |  |
| **AB <3 months** | yes | 81 | 22.6 | 777 | 29.4 | **0.007** |

Chi-square p-value for statistical significance of AOM vs DCC. The categorical variables were age (6-12, 13-24 and 25-30 moths), number of siblings (0, 1, >1) and a history of AOM (0, 1, >1). AOM= acute otitis media, DCC= day care centre, AB= antibiotics

The PCV10-vaccinated proportion increased over the study period from 2.6% to 83.9% and from 0.0% to 75.8%, in AOM children and DCC children respectively. The proportion of PCV13-vaccinated children decreased from 74.4% to 1.0% in AOM and 75.7% to 2.7% in DCCs (Figure 1). In each period the overall difference in vaccination status (as defined in figure 1) between AOM and DCC was significant.

**Figure 1:**
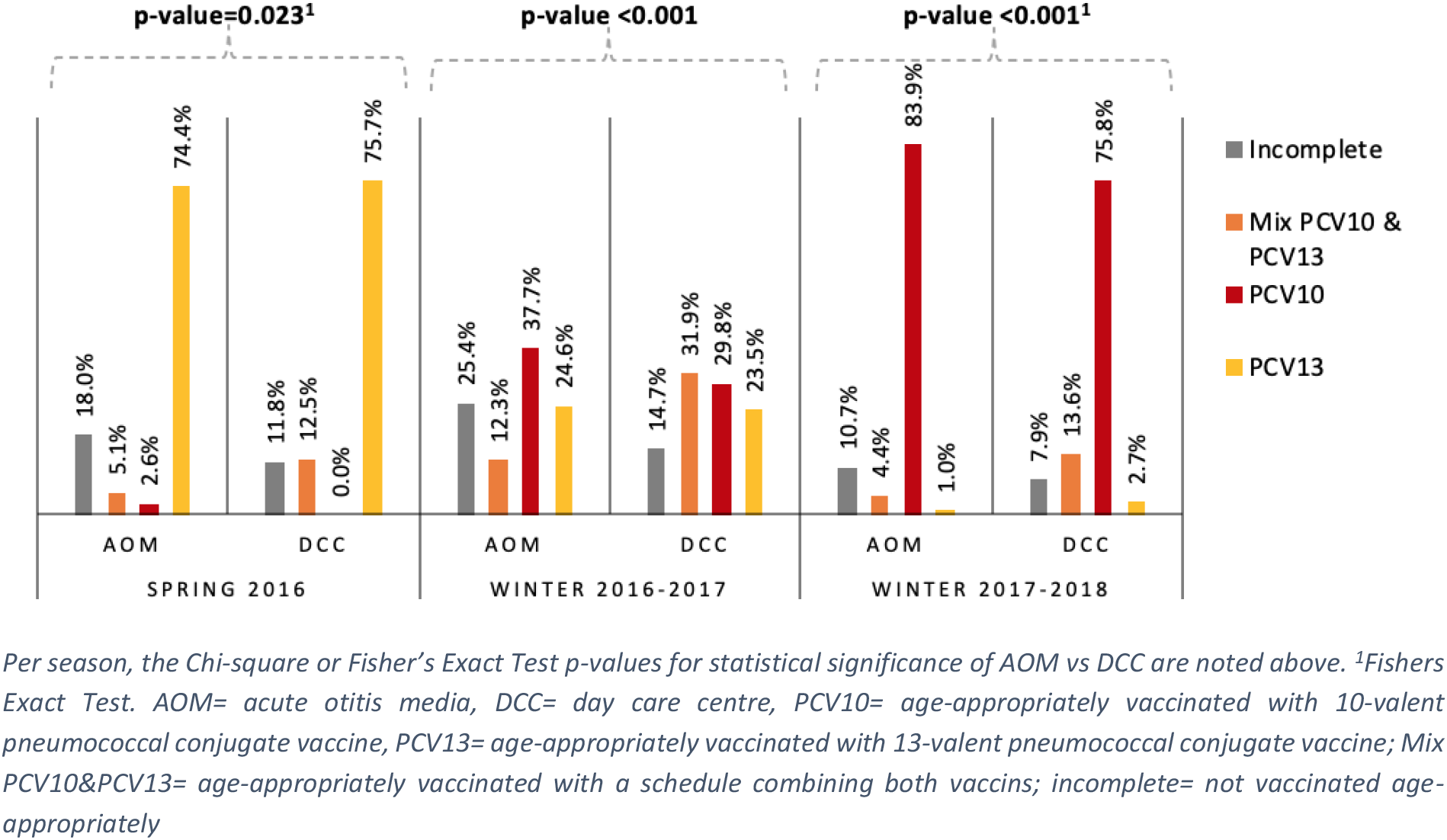
Vaccination status of children with AOM and children attending DCCs per season in Belgium in 2016-2018

### Carriage prevalence

Pneumococcal carriage prevalence was similar in AOM and in DCC, (AOM: 79.2%; DCC: 77.5%; p-value=0.454; Chi2) and among different age categories, for both AOM (6-12m: 76.7%, 13-24m: 82.5%, 25-30m: 77.8%; Chi2; p-value=0.431) and DCC (6-12m: 75.8%, 13-24m: 79.1%, 25-30m: 75.6%; Chi2; p-value=0.098).

### Serotype distribution

A total of 2241 Sp strains were cultured and 51 different serotypes identified, of which 34 serotypes in 268 strains from AOM children and 49 serotypes in 1973 strains from DCC children. In 65 DCC children and 7 AOM children, two serotypes were identified. PCV13 serotypes increased over time in AOM (Y1: 0.0%, Y2: 8.4%, Y3: 8.3%) and DCC (Y1: 1.8%, Y2: 2.6%, Y3: 9.8%), mainly because of an increased prevalence of serotype 19A (AOM: Y1: 0.0%, Y2: 3.6%, Y3: 6.4%; DCC: Y1: 0.6%, Y2: 1.9%, Y3: 8.8%). In children with AOM, PCV13 vaccine serotype 3 also increased in year 2 and year 3. Serotype 23B, a non-vaccine type, was most frequent in AOM (12.3%) and DCC (17.4%), followed by 11A (7.5%), 15B (7.5%) and 6C (6.3%) in AOM and 23A (8.0%), 11A (7.4%) and 15B (7.1%) in DCC. Serotypes 8 and 19B were found in AOM children (both 0.4%), but not in DCC children. When we ranked serotypes by frequency per study period and per study group, we found that the number of serotypes together accounting for 50% of carriage increased from 5 to 7 over time in both population groups (Table 8; Appendix 2). Among those dominating serotypes, we found 23B, 11A and 15 in all periods for both AOM and DCC and serotype 23A exclusively in DCC, whereas serotype 6C and 19A appeared in both groups in period 3. Frequency of vaccine serotypes over time was summarised separately in Table 9; Appendix 3. For 9 serotypes, frequency differed significantly between AOM and DCC (Table 3), although the difference was not significant in every separate period. Only serotype 17F showed a significant difference for more than one period: period 2 (AOM: 3.6%; DCC: 0.5%; FET; p-value=0.023) and period 3 (AOM: 2.6%; DCC: 0.6%; FET; p-value=0.043). The frequency of other serotypes, found either in AOM and DCC, was similar.

**Table 3:** Difference in serotype distribution of Streptococcus pneumoniae between children with AOM and children attending DCC in Belgium in 2016-2018.

|  | AOM frequency (%) | DCC frequency (%) | Statistical significance (p-value for AOM vs DCC) |
| --- | --- | --- | --- |
| **3** | 2.6 | 0.8 | 0.015<sup>1</sup> |
| **6C** | 6.3 | 3.4 | 0.030<sup>2</sup> |
| **7B** | 0.8 | 0.1 | 0.039<sup>1</sup> |
| **9N** | 2.2 | 0.9 | 0.048<sup>1</sup> |
| **12F** | 2.2 | 0.8 | 0.032<sup>1</sup> |
| **17F** | 2.6 | 0.6 | 0.003<sup>1</sup> |
| **23A** | 4.1 | 8.0 | 0.025<sup>2</sup> |
| **23B** | 12.3 | 17.4 | 0.036<sup>2</sup> |
| **29** | 1.1 | 0.2 | 0.041<sup>1</sup> |

Only serotypes that were found significantly more in one of the two groups are shown. <sup>1</sup>Fisher's Exact Test, <sup>2</sup>Chi2, AOM= acute otitis media, DCC= day care centre.

**Table 4:** Median Streptococcus pneumoniae carriage density, expressed in ×105 copies/μl, per age group for AOM and DCC in 2016-2018.

|  | 6-12 months | 13-24 months | 25-30 months |
| --- | --- | --- | --- |
| **AOM** | 3.90 | 4.78 | 9.47 |
| **DCC** | 2.53 | 5.20 | 3.33 |
| **Statistical significance (p-value MWU for AOM vs DCC)** | **0.035** | 0.819 | **0.045** |

AOM: 6-12 months N=117, 13-24 months N=115, 25-30 months N=30; DCC: 6-12 months N=366, 13-24 months N=1027, 25-30 months N=489). AOM= acute otitis media, DCC= day care centres, MWU= Mann-Whitney U Test.

### Carriage density of Sp positive samples

Overall, median pneumococcal DNA load was slightly higher, though non-significant, in AOM-infants (5.0 ×10^5^ copies/μl, 3.3 ×10^5^ – 7.6 ×10^5^ copies/μl) than in DCC-infants (4.2 ×10^5^ copies/μl, 3.8 ×10^5^ – 5.7 ×10^5^ copies/μl) (MWU for median; p-value=0.154). In 2017-2018, the year with the highest number of AOM-participants, median density in children with AOM almost doubled density in DCC children (AOM: 5.3 ×10^5^ copies/μl; DCC: 3.3 ×105 copies/μl; MWU p-value=0.086). In an age-specific analysis, median density was higher in carriers with AOM than in DCC in the 6-12 months and 25-30 months age category, but not in children aged 13-24 months .

### Antimicrobial non-susceptibility

Non-susceptibility of Sp to at least one antibiotic remained stable over the study period for both AOM (Y1: 48.2%, Y2: 51.9%, Y3: 48.1%; Chi^2^; p-value=0.851) and DCC (Y1: 47.0%, Y2: 48.4%, Y3: 44.6%; Chi^2^; p-value=0.352). Non-susceptibility to more than one antibiotic increased borderline non-significantly in the AOM population (Y1: 18.5%, Y2: 25.3%, Y3: 35.7%; Chi^2^; p-value=0.092) and increased significantly in the DCC population (Y1: 18.4%, Y2: 26.3%, Y3: 30.5%; Chi^2^; p-value<0.001). Lastly, 19A strains were more frequently resistant to penicillin in AOM children (AOM: 8.8%; DCC: 1.2%; FET: p-value=0.002).

### Clinical signs in children with AOM

Otoscopic signs are summarised in Table 5. Average temperature was 39.1°C (36.0°C - 41.6°C); in 15 children temperature was unknown. A small number of patients had no fever (8.7%, 30/347), 14.7% (51/347) had a mild fever of 38-38.5°C, 28.8% (100/347) had a fever of 38.6-39°C, while 47.8% (166/347) had a temperature of >39°C. Antibiotic prescription occurred in 77.3% of cases (279/361, 1 missing value), of which 91.0% (254/279) by a prescribing physician. Overall, 17.4% of AOM children (63/362) were seen by a non-prescribing physician.

**Table 5:** Clinical spectrum of acute otitis media. Missing values (otorrhea: 5, effusion: 10, bulging: 4) were due to incomplete questionnaires.

|  | None<br>% (n) | Unilateral<br>% (n) | Bilateral<br>% (n) |
| --- | --- | --- | --- |
| **Otorrhea** | 85.2 (304/357) | 11.2 (40/357) | 3.6 (13/357) |
| **Effusion** | 20.2 (71/352) | 33.8 (119/352) | 46.0 (162/352) |
| **Bulging** | 10.3 (37/358) | 71.0 (254/358) | 18.7 (67/358) |

A second visit for AOM within 1 month was reported for 65 children (18.0%), of whom 28 went to a physician participating in the study. However, not all parents could be reached for a follow-up phone call (24.6%, 89/362 missing). For six children antibiotics were prescribed in visit two, of whom five had also received antibiotics in the previous visit. Reasons for a second visit were: deterioration established by a physician (9.2%, 6/65), deterioration according to the parents during a follow-up phone call (10.8%, 7/65), additional infection established by a physician (24.6%, 16/65), relapse according to the parents during a follow-up phone call (35.4%, 23/65) or unknown reasons (20.0%, 13/65).

Median recovery time was four days; 47.5% of children recovered within three days (124/261), and 8.8% of children needed more than eight days (23/261). Missing values were due to no follow-up phone call (N=89) or incomplete phone call questionnaires (N=12). The only difference we found was AB use between children carrying vaccine serotypes and non-vaccine serotypes. Children with non-vaccine serotypes more often received antibiotics than children The clinical score data was left-skewed (more children with a lower clinical score). The median clinical score was 6/11, with a minimum and maximum score of 1/11 (5/356 children) and 10/11 (1/356 children), respectively. 44.9% of AOM episodes had a clinical score <6 and 55.1% had a higher score (Table 6), with no difference between non-vaccine and vaccine serotype carriers (FET; p-value=0.931; Table 6). Individual analysis per serotype was not possible.

**Table 6:** Clinical severity score in children with AOM per serotype group, Belgium 2016-2018. PCV7= 7-valent pneumococcal conjugate vaccine, PCV13= 13-valent pneumococcal conjugate vaccine

| capsulartype | Median clinical score | clinical score<6 |  | clinical score ≥6 |  |
| --- | --- | --- | --- | --- | --- |
|  |  | n | % | n | % |
| Non-vaccine type | 6 | 100 | 43.7 | 129 | 56.3 |
| PCV7 serotype | 6 | 3 | 37.5 | 5 | 62.5 |
| PCV13 serotype | 5.5 | 10 | 50.0 | 10 | 50.0 |
| Untypeable | 5 | 2 | 50.0 | 2 | 50.0 |

## Discussion

In this 3-year nasopharyngeal carriage study, differences were identified in S pneumoniaeserotype distribution but not in its antimicrobial resistance, between children with AOM and healthy children in DCC. No significant difference was found in overall Sp carriage prevalence between the two populations. This is plausible since AOM episode and DCC-attendance are both associated with high Sp carriage [19–22].

Serotypes 3, 6C, 9N, 12F, 17F, 7B and 29 were found more frequently in AOM children than in DCC, although they were all together uncommon (AOM: 17.9%; DCC: 6.7%). Serotype 3 was thus the only vaccine serotype that was identified significantly more in children with AOM than in children attending DCC which is in line with previous findings that associated serotype 3 to a high invasive capacity and more-severe disease [23–27]. Serotypes 9N and 12F have not previously been associated with AOM, though they have been associated with IPD: serotype 12F is frequent (20%) among infants with IPD in Belgium, and in Wales and England, serotypes 8, 12F and 9N increased rapidly and became responsible for more than 40% of total IPD cases after PCV13 introduction. [26, 40-43]. Serotype 17F has previously been associated with serious clinical outcomes and higher case-fatality, but not with AOM [44, 45]. Little is known about the invasive potential of Serotypes 7B and 29 since carriage of these serotypes is rare.

A similar study in Taiwan, investigated the serotype distribution of Sp in children with AOM and healthy children (control group) between the ages of 2-60 months after routine PCV13 immunization. In contrast to our findings, non-vaccine serotypes accounted for 71.4% of all serotyped isolates in the otitis group and 90.9% in the control group because the identified vaccine serotypes 19A (most common), 19F and 23F were more common in the AOM group than in the control group [46]. The most common serotypes in children with AOM as well as in healthy children in that study were 23A, 19A and 15B/C, whereas we found serotypes 23B and 11A and 15B very common in both AOM and DCC and serotype 23A in DCC children.

However, the prevalence of PCV13 serotypes increased over the three year study period in our study. Serotype 19A was the most frequent PCV13 serotype in period three for both populations and was found at similar frequency in the two groups in each period. In Belgian children, 19A was also the most frequently identified serotype in IPD (27%) and MEF (26%) in 2018 [28]. In other studies looking at clinical manifestation of AOM episodes by serotype, 19A was (one of) the most frequent serotype(s) identified [28, 29]. After PCV10 introduction in Bulgaria, the prevalence of 19A in the middle ear fluid, which is the most reliable to determine the causative serotype, of children diagnosed with AOM was 5.6% [30]. This is similar as the nasopharyngeal 19A carriage in we found in children with AOM in Belgium, two to three years after PCV10 introduction (6.4%). Chen et al. also found that 19A appears to have higher AOM potential, in comparison to other serotypes, while a nasopharyngeal carriage study in France found that children with AOM have a higher risk of 19A nasopharyngeal colonization [31, 32]. The PCV7 serotype 19F also continues to circulate and was found in children with AOM in all periods, as was the case in other PCV7- PCV10- and PCV13-administering countries [10, 27, 29, 33–37]. A Finnish Otitis Media Cohort Study found 19F was one of the most common serotypes isolated from middle ear infections in PCV7-vaccinated children [38, 39]. Among non-PCV13 serotypes, serotype 23B was most prevalent in each study population, as reported in other studies [10, 27, 34]. In this study, 23A and 23B were also found significantly more frequent in healthy children attending DCC, in comparison to children with AOM. Other studies showed that serotypes 23A and 23B are more commonly carried and carried for a longer period and can therefore circulate at a higher rate than other serotypes [25].

No significant difference in carriage density of Sp between children with AOM and children attending DCC was found. A higher density was expected in children with AOM, since a higher pneumococcal load is associated with disease [47–49]. However, looking at period three, the most representative year due to a higher number of children with AOM, the difference in median pneumococcal carriage density was borderline significant. high Sp density in the nasopharynx is suggested to be a predictor of transmission probability [50, 51] and of disease. Dunne et al. found that having upper respiratory tract infection symptoms was associated with an increased pneumococcal density [52–54]. In Belgium, 36.4% of children in DCCs presented common cold symptoms, which associates with the high pneumococcal density in DCC-children. Murad et al. demonstrated in regression analysis that density was lower in infants currently on antibiotics [52], but taking antibiotics was an exclusion criterium in both groups of children in our study. Fadlyana et al. found that low parental income (socioeconomic status) was associated with higher Sp density [51]. The background of parents is not known in this study, though children of a higher socioeconomic status tend to have more access to child care services, such as day care

Non-susceptibility to one or more antibiotics was similar in Sp strains found in children with AOM and children attending DCC and was within the range identified in other PCV-using countries [10, 30, 33, 55, 56]. ST19A has been reported as a serotype that is frequently non-susceptible to antibiotics, especially after PCV7 or PCV10 introduction [10, 57, 58]. However, 19A penicillin non-susceptibility was still at the lower end of the range (AOM: 8.8%; DCC: 1.2%). After PCV7 introduction in France, penicillin non-susceptibility for serotype 19A in children with AOM ranged from 77.6-92.0% [31]. Since children who were treated with oral ABs in the seven days prior to sampling were excluded, some non-susceptible strains could have been missed.

PCV introduction was reported to reduce AOM-related doctor visits as well as risk of early first AOM episode [5, 59–63]. The higher risk for children with an incomplete vaccination status, in comparison to fully PCV13-vaccinated children, for having AOM is in line with these findings.

Finally, for 25-to-30-month-olds, a three times higher carriage density was observed in the AOM-population compared to the DCC-population. Since risk of AOM development decreases with age [64], this finding suggests a role for carriage density in disease development.

Only few existing studies are comparing the clinical picture of AOM caused by different serotypes [29, 30, 65]. The strength of the current study is that symptoms are not interpreted by the child’s parents or caregivers, introducing bias, but by objective interpretation of symptoms by a pediatrician or doctor. In the current study, serotypes 16F (N=6), 24B (N=3) and 12F (N=6) were associated with the highest clinical scores. Previous studies associated serotype 16F mainly with asymptomatic carriage [66, 67]. For all 12F AOM episodes, antibiotics were prescribed. This is in line with previous findings that associated serotype 12F with severe (invasive) disease [42, 66]. Serotype 23B had also a high clinical score (≥6) in more than 70% of AOM episodes and antibiotics were prescribed for 93.6% of episodes (overall 77.5%). This is possibly problematic, since serotype 23B was also most often identified as cotrimoxazole (AOM: 27.8%; DCC: 31.8%) and penicillin (AOM: 36.8%; DCC: 37.7%) non-susceptible in this study. Another serotype with a high clinical score (in more than 70% of AOM episodes) was the vaccine serotype 19F (included in PCV7, PCV10 and PCV13), which also had the highest frequency of second visits (42.9%). Serotype 19F has been associated with invasive infections in children and AOM with spontaneous otorrhea [24, 32, 68, 69]. Furthermore, Palmu et al. found serotype 19F as one of the most common pneumococcal serotypes recovered from samples of MEF and the serotype most often leading to earache [68, 70]. In our study, carried 19F also had a high rate of resistance to at least one antibiotic (85.7%).

A major strength of this study is the combined use of culture and molecular methods to explore pneumococcal carriage in nasopharyngeal microbiota. PCR is more sensitive to detect carriage and easier to quantify, however, standard culture methods still allow the most extensive and reliable serotype discrimination and antibiotic susceptibility assessment. However, some limitations should be considered when interpreting the presented results. First, in children with AOM we cannot be sure that the carried Sp serotype was the cause of the AOM episode, especially since 91.4% and 95.5% of Sp positive samples also carried *Haemophilus influenzae* and *Moraxella catarrhalis.* Middle ear fluid samples allow for more accurate identification of the causative agent, but are not standard practice in first-line AOM approach in Belgium. Second, findings refer to the specific child populations that were studied and cannot be generalized to nonresponsive or chronic otitis media, to children of all ages, or even to all AOM episodes. Moreover, children attending day care might share more characteristics with children with AOM than the healthy child population in general [71] limiting the power to detect any differences. In addition, 148 DCC children representing 303 isolates participated more than once, but in different study periods. A sensitivity analysis that only used their first sample confirmed all findings for child characteristics, carriage prevalence and carriage density. Finally, any trends in time should be interpreted cautiously. Study population characteristics were not constant over time, especially in the AOM population in whom recruitment was more successful towards the end of the study. Moreover, calendar time was different in the first collection period than in the later ones, which might have exaggerated season-dependent changes in serotype distribution. However, the marked 19A increase was the only time trend we identified, a trend we suppose not to be due to differences in season as no season-dependency of 19A carriage has been reported so far. Still, natural fluctuation cannot be excluded since the study period is quite short.

## Conclusion

This large surveillance study in Belgium monitored pneumococcal carriage in the nasopharynx of healthy children in day care and children with AOM. Overall carriage prevalence and density were similar for the two child populations. The predominant serotypes after PCV10 implementation were 23B, 23A, 11A and 15B, of which almost half of carried strains showed non-susceptibility to one (or more) antibiotics. Serotypes 3, 6C, 9N, 12F, 17F, 7B and 29 were less common but were found more frequently in AOM children than in DCC. The change in vaccination program induced similar changes in both populations, with an increase of PCV13-serotype 19A over the study period from 0.0% in year 1 to 6.4% in year 3 for the AOM-group and 0.6% in year 1 to 8.8% in year 3 for the DCC-group. We can conclude that young children with AOM did not carry Sp more frequently or at higher load than healthy children in DCC, but the serotypes that were more frequently found in AOM were mainly non-vaccine types. The continued surveillance of Sp carriage in children attending DCC will demonstrate whether the recent switch from PCV10 back to PCV13 in the infant programme in Belgium will further change the carriage of Sp.

## Acknowledgements

The authors would like to thank the members of the expert advisory board for their contribution to the interpretation of the results; Research Link–ECSOR operating as the CRO; the cooperating nurses, physicians, the participating baby clinics for assistance in the recruitment and sampling; the children and their parents for their participation.

## NPcarriage Study Group

David Tuerlinckx, CHU Dinant-Godinne, Université Catholique de Louvain, Yvoir, Belgium; Adam Finn, University of Bristol, School of Cellular and Molecular Medicine, Bristol, UK; Koen Van Herck, Department of Public Health and Primary Care, Ghent University, Gent, Belgium, Centre for the Evaluation of Vaccination, Vaccine and Infectious Disease Institute, University of Antwerp, Wilrijk, Belgium; Robert Cohen, Université Paris Est, IMRB-GRC GEMINI, 94000 Créteil, France; Christine Lammens, Laboratory of Medical Microbiology, Vaccine and Infectious Disease Institute, University of Antwerp, Wilrijk, Belgium; Marc Verghote (Private pediatrician) ; Marc De Meulemeester (Private pediatrician); Barbara De Wilde (Private pediatrician); Kate Sauer, Pediatric Department, AZ Sint-Jan, Brugge, Belgium; Luc Pattyn, Pediatric Department, AZ Turnhout, Belgium; Bart Rutteman, Pediatric Department, AZ Sint-Blasius, Dendermonde, Belgium; Caroline Genin, Pediatric Department, C.H.C Espérance, Saint-Nicolas, Belgium; Diane Stroobant, Pediatric Department, Grand Hôpital de Charleroi, Charleroi, Belgium; Ilse Ryckaert, Pediatric Department AZ-Nikolaas, Sint-Niklaas, Belgium; Tessa Goetghebuer, Pediatric Department, CHU Saint-Pierre, Brussels, Belgium; Zornitsa Vassileva, Pediatric Department, AZ Lokeren, Lokeren, Belgium

## Appendices

## Appendix 1

**Table 7:** Demographic and clinical characteristics of children with AOM per period with Chi-square p-value for trend in Belgium in 2016-2018.

|  |  | Period 1:<br>2016<br>N=39 |  | Period 2:<br>2016-2017<br>N=122 |  | Period 3:<br>2017-2018<br>N=205 |  | P-value<br>Chi<sup>2</sup> for trend |
| --- | --- | --- | --- | --- | --- | --- | --- | --- |
|  |  | n | % | n | % | n | % |  |
| Region | Flanders | 8 | 20.5 | 70 | 57.4 | 119 | 58.1 | <0.001 |
|  | Wallonia | 27 | 69.2 | 46 | 37.7 | 68 | 33.2 |  |
|  | Brussels | 4 | 10.3 | 6 | 4.9 | 18 | 8.8 |  |
| Age (months) | 6-12 | 14 | 35.9 | 57 | 48.3 | 92 | 44.9 | 0.716 |
|  | 13-24 | 19 | 48.7 | 46 | 39.0 | 89 | 43.4 |  |
|  | 25-30 | 6 | 15.4 | 15 | 12.7 | 24 | 11.7 |  |
| Gender | male | 20 | 51.3 | 64 | 54.2 | 102 | 49.8 | 0.740 |
| Preterm delivery | yes | 0 | 0.0 | 13 | 11.2 | 12 | 5.9 | 0.038 |
| Breastfeeding >6 months | yes | 7 | 18.0 | 26 | 22.4 | 48 | 23.4 | 0.755 |
| At least one parent smokes | yes | 15 | 38.5 | 29 | 25.0 | 59 | 28.8 | 0.273 |
| Previous hospitalisation | yes | 13 | 33.3 | 33 | 28.5 | 62 | 30.2 | 0.842 |
| Number of siblings | 0 | 3 | 7.7 | 30 | 26.1 | 84 | 41.0 | <0.001 |
|  | 1 | 14 | 35.9 | 41 | 35.7 | 66 | 32.2 |  |
|  | >1 | 22 | 56.4 | 44 | 38.2 | 55 | 26.8 |  |
| AOM-history | 0 | 0 | 0.0 | 0 | 0.0 | 101 | 49.3 | <0.001 |
|  | 1 | 13 | 52.0 | 22 | 41.5 | 51 | 24.9 |  |
|  | >1 | 12 | 48.0 | 31 | 58.5 | 53 | 25.9 |  |
| AB <3 months | yes | 11 | 28.2 | 27 | 23.5 | 43 | 21.0 | 0.588 |

AOM= acute otitis media, AB= antibiotics.

## Appendix 2

**Table 8:**
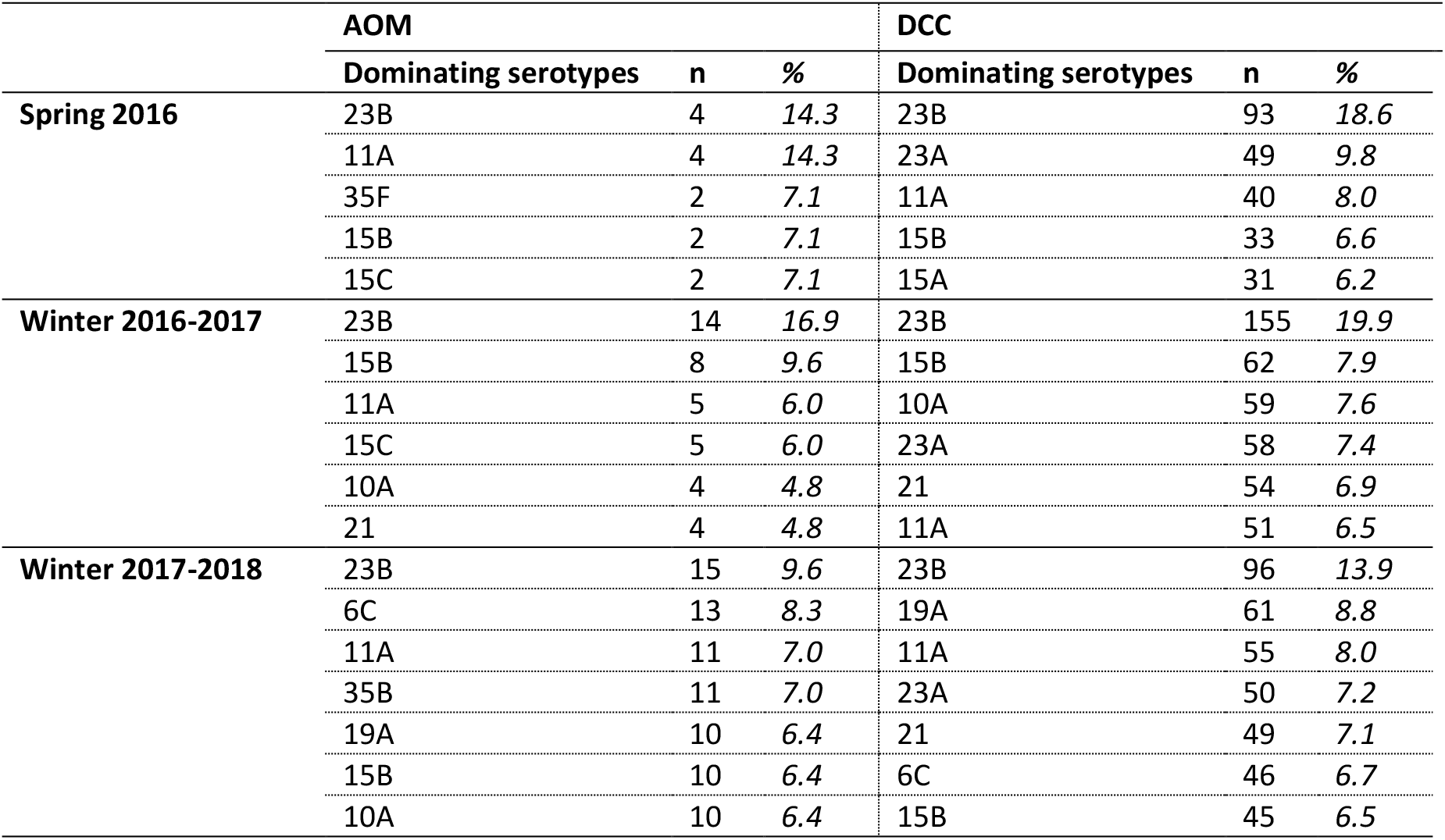
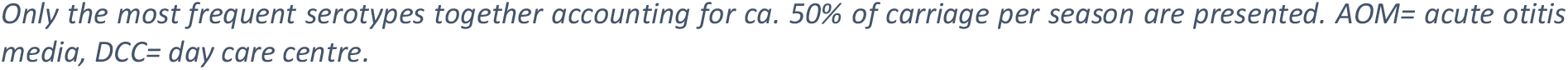
Carriage frequency of Streptococcus pneumoniae serotypes in the AOM child population and the healthy child population attending DCC per season in Belgium in 2016-2018.

|  | AOM |  |  | DCC |  |  |
| --- | --- | --- | --- | --- | --- | --- |
|  | Dominating serotypes | n | % | Dominating serotypes | n | % |
| Spring 2016 | 23B | 4 | 14.3 | 23B | 93 | 18.6 |
|  | 11A | 4 | 14.3 | 23A | 49 | 9.8 |
|  | 35F | 2 | 7.1 | 11A | 40 | 8.0 |
|  | 15B | 2 | 7.1 | 15B | 33 | 6.6 |
|  | 15C | 2 | 7.1 | 15A | 31 | 6.2 |
| Winter 2016-2017 | 23B | 14 | 16.9 | 23B | 155 | 19.9 |
|  | 15B | 8 | 9.6 | 15B | 62 | 7.9 |
|  | 11A | 5 | 6.0 | 10A | 59 | 7.6 |
|  | 15C | 5 | 6.0 | 23A | 58 | 7.4 |
|  | 10A | 4 | 4.8 | 21 | 54 | 6.9 |
|  | 21 | 4 | 4.8 | 11A | 51 | 6.5 |
| Winter 2017-2018 | 23B | 15 | 9.6 | 23B | 96 | 13.9 |
|  | 6C | 13 | 8.3 | 19A | 61 | 8.8 |
|  | 11A | 11 | 7.0 | 11A | 55 | 8.0 |
|  | 35B | 11 | 7.0 | 23A | 50 | 7.2 |
|  | 19A | 10 | 6.4 | 21 | 49 | 7.1 |
|  | 15B | 10 | 6.4 | 6C | 46 | 6.7 |
|  | 10A | 10 | 6.4 | 15B | 45 | 6.5 |

Only the most frequent serotypes together accounting for ca. 50% of carriage per season are presented. AOM= acute otitis media, DCC= day care centre.

## Appendix 3

**Table 9:** Prevalence of S. pneumoniae serotypes overall and per season in Belgium in 2016-2018.

|  |  | General |  | Spring 2016 |  | Winter 2016-2017 |  | Winter 2017-2018 |  |
| --- | --- | --- | --- | --- | --- | --- | --- | --- | --- |
|  |  | AOM<br>n (%) | DCC<br>n (%) | AOM<br>n (%) | DCC<br>n (%) | AOM<br>n (%) | DCC<br>n (%) | AOM<br>n (%) | DCC<br>n (%) |
| **PCV7 VT**<br>**prevalence (%)** | **19F** | 7 (2.6) | 42 (2.1) | 1 (3.6) | 13 (2.6) | 2 (2.4) | 13 (1.7) | 4 (2.6) | 16 (2.3) |
|  | **23F** | 0 (0.0) | 3 (0.2) | 0 (0.0) | 3 (0.6) | 0 (0.0) | 0 (0.0) | 0 (0.0) | 0 (0.0) |
|  | **14** | 1 (0.4) | 5 (0.3) | 1 (3.6) | 4 (0.8) | 0 (0.0) | 1 (0.1) | 0 (0.0) | 0 (0.0) |
| **PCV10 VT**<br>**prevalence (%)** | **1** | 0 (0.0) | 1 (0.1) | 0 (0.0) | 1 (0.2) | 0 (0.0) | 0 (0.0) | 0 (0.0) | 0 (0.0) |
| **PCV13 VT**<br>**prevalence (%)** | **19A** | 13 (4.9) | 79 (4.0) | 0 (0.0) | 3 (0.6) | 3 (3.6) | 15 (1.9) | 10 (6.4) | 61 (8.8) |
|  | **3** | 7 (2.6) | 16 (0.8) | 0 (0.0) | 4 (0.8) | 4 (4.8) | 5 (0.6) | 3 (1.9) | 7 (1.0) |
|  | **6A** | 0 (0.0) | 2 (0.1) | 0 (0.0) | 2 (0.4) | 0 (0.0) | 0 (0.0) | 0 (0.0) | 0 (0.0) |
| **NVT prevalence**<br>**(%)** | **23B** | 33 (12.3) | 344 (17.4) | 4 (14.3) | 93 (18.6) | 14 (16.9) | 155 (19.9) | 15 (9.6) | 96 (13.9) |
|  | **11A** | 20 (7.5) | 146 (7.4) | 4 (14.3) | 40 (8.0) | 5 (6.0) | 51 (6.5) | 11 (7.0) | 55 (8.0) |
|  | **15B** | 20 (7.5) | 140 (7.1) | 2 (7.1) | 33 (6.6) | 8 (9.6) | 62 (7.9) | 10 (6.4) | 45 (6.5) |
|  | **6C** | 17 (6.3) | 68 (3.5) | 1 (3.6) | 9 (1.8) | 3 (3.6) | 13 (1.7) | 13 (8.3) | 46 (6.7) |
|  | **10A** | 15 (5.6) | 111 (5.6) | 1 (3.6) | 27 (5.4) | 4 (4.8) | 59 (7.6) | 10 (6.4) | 25 (3.6) |
|  | **35B** | 15 (5.6) | 94 (4.8) | 1 (3.6) | 17 (3.4) | 3 (3.6) | 42 (5.4) | 11 (7.0) | 35 (5.1) |
|  | **35F** | 12 (4.5) | 73 (3.7) | 2 (7.1) | 14 (2.8) | 2 (2.4) | 35 (4.5) | 8 (5.1) | 24 (3.5) |
|  | **23A** | 11 (4.1) | 157 (8.0) | 1 (3.6) | 49 (9.8) | 3 (3.6) | 58 (7.4) | 7 (4.5) | 50 (7.2) |
|  | **21** | 11 (4.1) | 126 (6.4) | 0 (0.0) | 23 (4.6) | 4 (4.8) | 54 (6.9) | 7 (4.5) | 49 (7.1) |
|  | **15C** | 11 (4.1) | 78 (4.0) | 2 (7.1) | 18 (3.6) | 5 (6.0) | 36 (4.6) | 4 (2.6) | 24 (3.5) |
|  | **15A** | 10 (3.7) | 95 (4.8) | 1 (3.6) | 31 (6.2) | 2 (2.4) | 41 (5.3) | 7 (4.5) | 23 (3.3) |

AOM= acute otitis media, DCC= day care centre, PCV7= 7-valent pneumococcal conjugate vaccine PCV10= 10-valent pneumococcal conjugate vaccine, PCV13= 13-valent pneumococcal conjugate vaccine, NVT= non-vaccine serotype.

